# Comparing the selective landscape of TLR7 and TLR8 across primates reveals unique sites under positive selection in *Alouatta*

**DOI:** 10.1101/708008

**Authors:** Nicole S. Torosin, Hernan Argibay, Timothy H. Webster, Patrice Showers Corneli, Leslie A. Knapp

**Affiliations:** Department of Anthropology, University of Utah, 260 S. Central Campus Dr., Salt Lake City, UT 84112; Department of Biology, University of Utah, 257 S. 1400 E., Salt Lake City, UT 84112; Instituto de Ecología, Genética y Evolución de Buenos Aires (IEGEBA-CONICET), Intendente Güiraldes 2160 - Ciudad Universitaria (C1428EGA) Ciudad Autónoma de Buenos Aires, Argentina

## Abstract

Among primates, susceptibility to yellow fever (YFV), a single-stranded (ss) RNA virus, ranges from complete resistance to high susceptibility. Howler monkeys (genus *Alouatta*) are the most susceptible to YFV. In order to identify *Alouatta*-specific genetic factors that may be responsible for their susceptibility, we collected skin samples from howler monkey museum specimens of the species *A. caraya* and *A. guariba clamitans*. We compared the rate of nonsynonymous to synonymous (*dN/dS*) changes of Toll-like receptor (TLR) 7 and TLR8, the two genes responsible for detecting all ssRNA viruses, across the Primate order. Overall, we found that the TLR7 gene is under stronger purifying selection in howler monkeys compared to other New World and Old World primates, but TLR8 is under the same selective pressure. When we evaluated *dN/dS* at each codon, we found six codons under positive selection in *Alouatta* TLR8 and two codons under positive selection in TLR7. The changes in TLR7 are unique to *A. guariba clamitans* and are found in functionally important regions likely to affect detection of ssRNA viruses by TLR7/TLR8, as well as downstream signaling. These amino acid differences in *A. guariba clamitans* may play a role in YFV susceptibility. These results have implications for identifying genetic factors affecting YFV susceptibility in primates.

## 1 Introduction

Host-pathogen interactions are major drivers of immune gene evolution (Sironi et al., 2015). Viruses, in particular, have caused close to 30% of all adaptive amino acid changes in mammals (Enard et al., 2016). Genetic comparisons between species can help identify substitutions differentially affecting disease susceptibility or resistance (Wlasiuk and Nachman, 2010; Areal et al., 2011; Schad and Voigt, 2016).

Among primates, susceptibility to yellow fever virus (YFV) ranges from complete resistance to high susceptibility (**Table 1**). Howler monkeys (genus *Alouatta*) are the most susceptible and usually die within a week of infection (Bicca-marques and Freitas, 2010). Currently, there is no explanation for *Alouatta’s* high level of susceptibility to YFV.

**Table 1:**
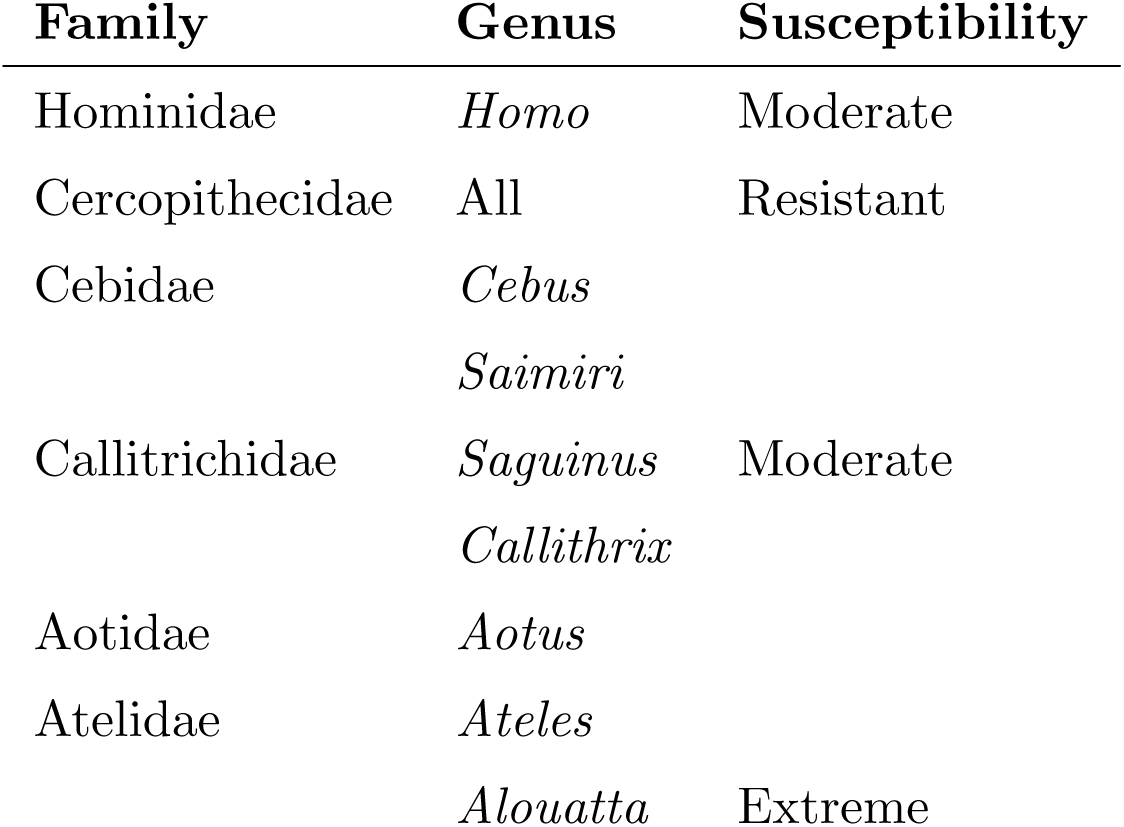
Primate susceptibility to YFV, listed by increasing susceptibility (Bicca-marques and Freitas, 2010).

In general, to understand species-specific genetic changes in response to pathogens, researchers compare the selective landscape of immune genes (Quach et al., 2013; Wlasiuk and Nachman, 2010). TLRs, such as TLR7 and TLR8, are evolutionarily-constrained innate immune genes specifically responsible for detecting single-stranded (ss) RNA viruses, such as West Nile virus, dengue virus, and YFV (Kawai and Akira, 2010; Kumar et al., 2011; Hanley et al., 2013). When these two genes detect ssRNAs, they initiate a signaling cascade that prompts the immune system to destroy infected cells and activate adaptive immunity (Kumar et al., 2011).

TLR7 and TLR8 are under strong purifying selection in humans and great apes (Quach et al., 2013). Amino acid changes that do occur are likely shaped by environmental or species-specific pathogens (Wlasiuk and Nachman, 2010). Polymorphisms within TLR7 and TLR8 have been implicated in progression of various human diseases such as Hepatitis C and Congo hemorrhagic fever (Lin et al., 2012). Therefore, TLR7 and TLR8 are good candidates to study variable susceptibility to ssRNA viral pathogens.

However, nothing is known about evolutionary patterns of endosomal TLRs in *Alouatta* and other New World primates. To address the question of *Alouatta’s* susceptibility to YFV, we collected skin samples from one taxidermied howler monkey of each species, *A. caraya* and *A. guariba clamitans*, and compared the selective landscape of newly sequenced TLR7 and TLR8 exons from *A. caraya* and *A. guariba clamitans* to TLR7 and TLR8 from published New World and Old World primates less susceptible to YFV (**Table 2**). Our comparison of TLR7 and TLR8 across primate species will help identify amino acid substitutions and selection patterns that may contribute to the high susceptibility of *Alouatta* to YFV.

**Table 2:**
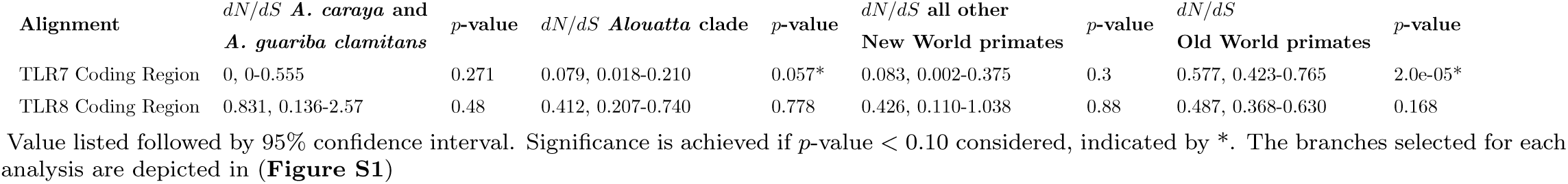
Substitution rates for tree partitions.

## 2 Materials and Methods

### 2.1 Sample collection and processing

Initially, we collected one *Alouatta caraya* liver sample from an individual that died during the 2007-2009 YFV outbreak in Misiones, Argentina as part of a YFV epizootic research project conducted by WCS-Global Health Program (**Table S1**). We stored the tissue at −70°*C* in 100% ethanol. We also collected 1 − 1.5 cm skin samples from one taxidermied *A. caraya* and one *A. guariba clamitans* museum specimen at the Museo Argentina de Ciencias Naturales Bernardino Rivadavia. We stored each skin sample in a sterile 1.5 ml Eppendorf tube at ambient temperature until DNA extraction.

We transported the tissue and taxidermied skin samples to the Universidad de Misiones at Posadas, Misiones, Argentina (transport permit 005436 issued by the “Jefatura de Gabinete de Ministros Secretaría de Ambiente y Desarrollo Sustentable”). At the Universidad, we extracted DNA using a Qiagen Fast DNA tissue kit (Qiagen, GmbH, Germany). We transported extracted DNA to Buenos Aires (transport permit 001494 issued by the “Presidencia de la Nación Secretaría de Ambiente y Desarrolo Sustentable”). We exported extracted DNA from Argentina using CITES export permit 042117 and CDC import permit number 2017-03-014.

To obtain DNA sequences, the University of Utah Huntsman Cancer Institute High-Throughput Genomics Center prepared and shotgun sequenced the *A. caraya* liver tissue DNA on an Illumina HiSeq 2500. The coding regions of TLR7 and TLR8 are highly conserved among primate species (Wlasiuk and Nachman, 2010), so we used BWA-MEM (Li, 2013) to align the reads from the three exons of TLR7 and two exons of TLR8 to a human reference sequence (1000 Genomes Project Consortium et al., 2015) and an unpublished *A. palliata* reference sequence (A. Burrell, pers. comm.) to confirm alignment. We aligned mtDNA genes ND1, ND2, and CO1 to the *A. palliata* reference sequence. We used SPAdes software to create an *A. caraya* TLR7 and TLR8 consensus sequence (Bankevich et al., 2012) (**Appendix: Methods**).

Using the newly generated *A. caraya* TLR7 and TLR8 reference sequences, we created a custom Ampliseq library IAD149391-168 (LifeTechnologies, Austin, TX). The University of Utah Sequencing Core completed targeted sequencing of the *A. caraya* and *A. guariba clamitans* skin samples using an Ion PGM and standard library kit protocols. To process the sequencing reads, we first trimmed the low quality bases from the ends and filtered fastq reads to maq > 20 using BBDuk (http://jgi.doe.gov/data-and-tools/bb-tools/). Next we aligned trimmed fastq reads for each sample to the *A. caraya* TLR7 and TLR8 reference sequences using BWA-MEM (Li, 2013). We used Freebayes (Garrison and Marth, 2012) to call variants for each sample. We used vcftools to remove indels and filter all sites with GQ *<* 30 (Danecek et al., 2011). We used bcftools to create a consensus sequence for each taxidermied skin sample (Li et al., 2009) (**Appendix: Methods**). GenBank accession numbers for all sequences generated can be found in **Table S2**.

DNA damage results in deamination of cytosine and guanine resulting in and excess of tyrosine and adenine substitutions, especially at the ends of sequencing reads (Briggs et al., 2007). Museum tissue samples are old and often subject to DNA damage, therefore we measured DNA damage using mapDamage2.0 software (Jónsson et al., 2013).

### 2.2 Published data and sequence alignment

We obtained coding regions of TLR7 and TLR8 for New World and Old World primates from GenBank and dnazoo.org (Dudchenko et al., 2017) (**Table S3**). We aligned DNA sequences using MEGA7 (Kumar et al., 2016) and confirmed amino acid translation of TLR7 and TLR8 using protein reference NP 057646.1 and NP 619542.1, respectively. We aligned and edited TLR7 and TLR8 amino acid translations for all species using Aliview (Larsson, 2014).

### 2.3 Phylogenetic analyses

Optimal phylogenetic models for each gene were chosen on the basis of the Decision-Theoretic (DT) method (Sullivan and Joyce, 2005) as implemented in PAUP 4.0 (Swofford, 2003) for model selection and estimating parameters (**Table S4**). We optimized the trees using these models in the Seaview implementation of PhyML (Guindon et al., 2010). Approximate likelihood ratio tests (aLRT) assign clade support levels across the tree (Anisimova and Gascuel, 2006).

### 2.4 Maximum Likelihood Selection Analysis

To test the hypothesis that the *Alouatta* lineage evolved under a different selection regime from other primates, we used the the positive selection option in HyPhy to “test whether a branch (or branches) in the tree evolve under different *dN/dS* than the rest of the tree” (HyPhy software (Kosakovsky Pond et al., 2005)). This analysis assumes a Muse Gaut codon model and is designed to test whether the ratio of non-synonymous to synonymous (*dN/dS*) substitutions is equal across the phylogenetic tree or is best described by multiple rates (Frost et al., 2005; Muse and Gaut, 1994). We compared selection coefficients of the following bipartitions: 1) the branch leading to newly sequenced species *A. guariba clamitans* and *A. caraya* only; 2) the branch leading to the whole *Alouatta* genus; and 3) the branch leading to all other New World primates (**Figure S1**).

A Fixed Effect Likelihood (FEL) test (HyPhy software (Kosakovsky Pond et al., 2005)) identified specific codons under selection in the *Alouatta* lineage. The null model assumes that the selection pressure for each site is constant for the entire alignment. The alternative model allows the *dN/dS* rate to vary site by site along the alignment. Results indicate whether certain sites in a selected lineage have been subject to positive or purifying selection. This analysis is designed for small sample sets and minimizes false positives (Kosakovsky Pond and Frost, 2005). We also completed FEL analysis for all other New World primates and Old World primates to see whether sites under selection in *Alouatta* are unique to the lineage.

### 2.5 Functional assessment

We assessed codon sites that showed evidence of positive selection for functional purpose. The TLR7 ligand binding region is within codons 500-589 in protein reference NP 057646.1 and within codons 481-577 in protein reference NP 619542.1 for TLR8 (Wei et al., 2009). Leucine-rich repeats (LRR), non-LRR insertions, and the Toll-interleukin receptor (TIR) domain for TLR7 and TLR8 are outlined in Bell et al. (2003). We examined codon sites in LRR, non-LRR insertions, and TIR domains (Bell et al., 2003) to determine if there was evidence for positive selection within any of the binding, LRR, insertion, or TIR regions.

## 3 Results

### 3.1 Sequences

We generated two *A. caraya* sequences and one *A. guariba clamitans* sequence for nuclear genes TLR7, exons 1 (221 bp), 2 (196 bp), and 3 (5635 bp); TLR8, exons 1 (201 bp) and 2 (4217 bp); and mtDNA genes ND1 (1075 bp), ND2 (1145 bp), and CO1 (1742 bp). Average sequencing coverage is reported in **Table S5, Table S6**. We did not detect post-mortem DNA damage in the museum samples (**Table S7**).

### 3.2 Phylogenetic trees

Maximum likelihood phylogenetic trees (**Figure 1, Figure 2**) show *Alouatta* species clustering next to other New World primates. The separate clustering of *Alouatta* is supported by other studies using nuclear genes (Steiper and Ruvolo, 2003; Boissinot et al., 1998; Harada et al., 1995) and mtDNA (**Figure S2**). Old World primates cluster as anticipated (Pozzi et al., 2014). In both nuclear DNA trees, the *A. guariba clamitans* branch length is longer than the other two *Alouatta* species, and in TLR7, it is the longest in the entire tree (**Figure 1, Figure 2**).

**Figure 1:**
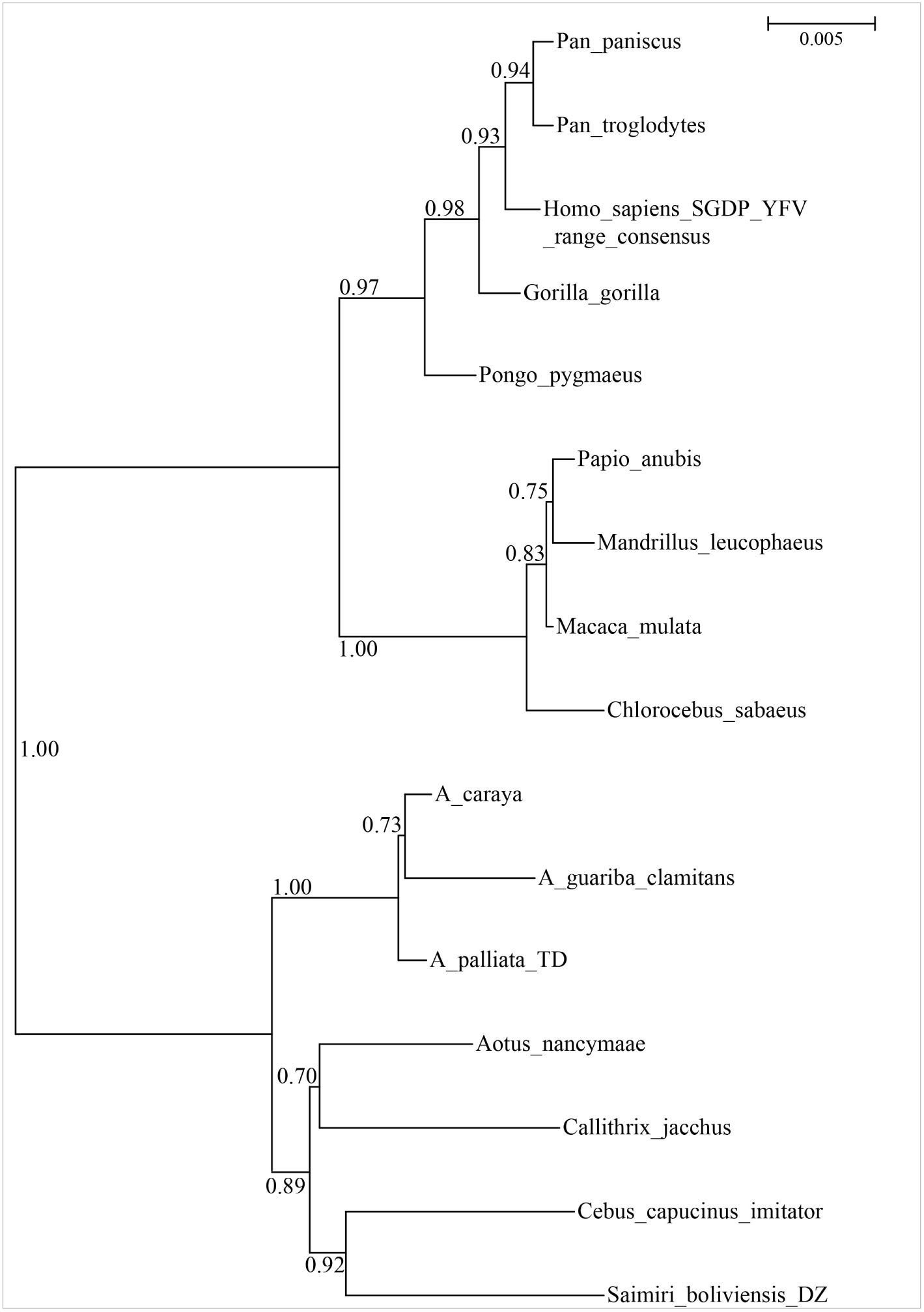
TLR7 maximum likelihood phylogenetic tree. TLR7 maximum likelihood phylogenetic tree generated from *A. caraya* and *A. guariba clamitans* TLR7 sequences (this study), New World and Old World primate TLR7 sequences. Each sequence totaled 3141 base pairs equaling 1039 codons. HKY model used, chosen by Decision Theoretic method. Log likelihood of the tree = -6740.7 and aLRT support values on branches.

**Figure 2:**
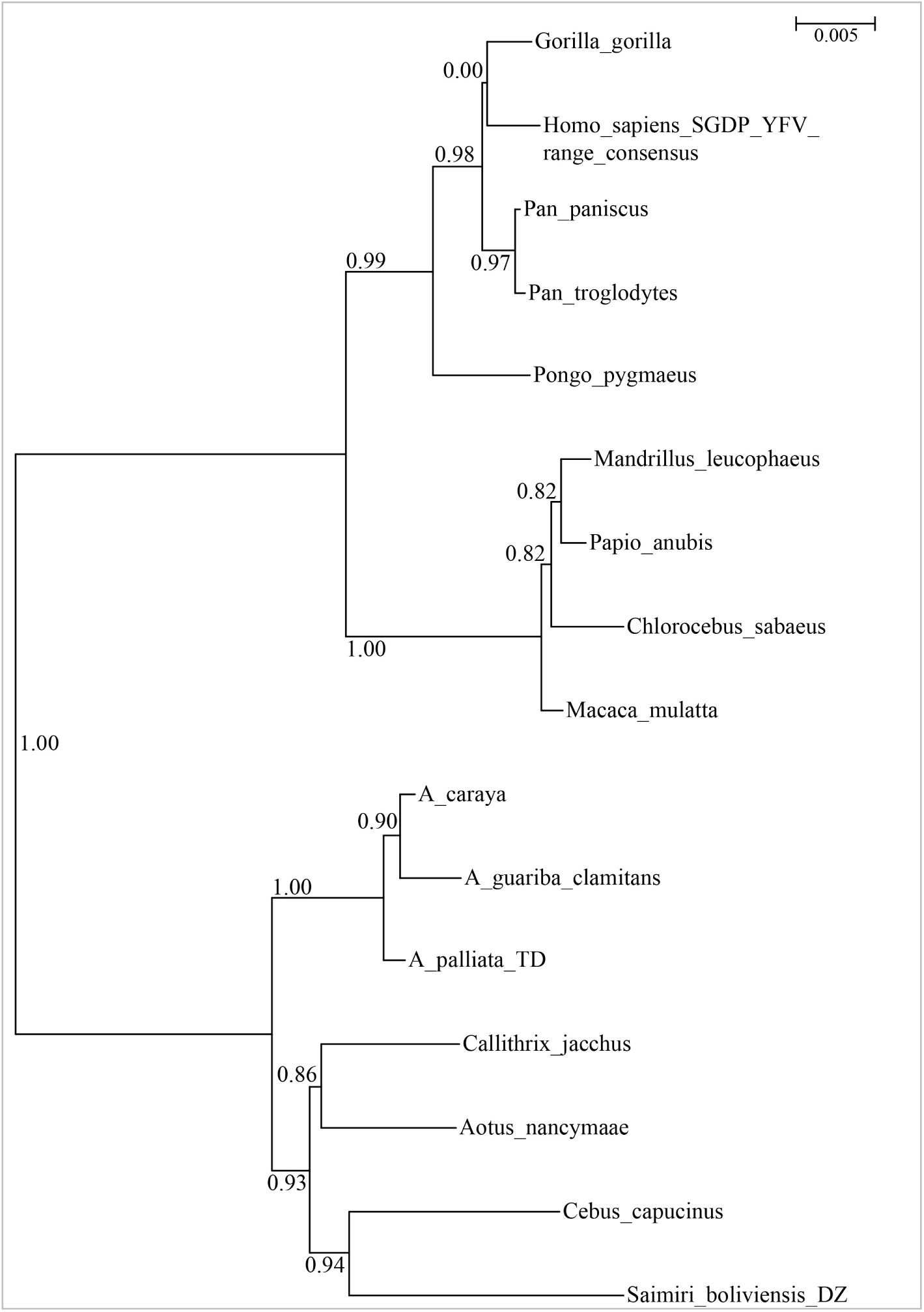
TLR8 maximum likelihood phylogenetic tree. TLR8 maximum likelihood phylogenetic tree generated from *A. caraya* and *A. guariba clamitans* TLR8 sequences (this study), New World and Old World primate TLR8 sequences. Each sequence totaled 3117 base pairs equaling 1047 codons. TN93 model used, chosen by Decision Theoretic method. Log likelihood of the tree = -7145.9 and aLRT support values on branches.

### 3.3 Substitution rates

We used a maximum likelihood approach to determine whether *Alouatta* branches evolved under a different *dN/dS* ratio from the rest of the primate tree. For this test, significance is reached at 0.10 because the standard 0.05 cutoff is too conservative and eliminates too many observations that deviate from the null hypothesis (Kosakovsky Pond and Frost, 2005). Our null hypothesis was that *dN/dS* is the same across the entire phylogenetic tree (Frost et al., 2005). We compared the following tree partitions: branches leading to the *A. caraya* and *A. guariba clamitans* clade, the branch leading to the whole *Alouatta* clade, the branch leading to the other New World primates, as well as the branch leading to the Old World primates (see **Figure S1** for illustration of branches tested).

Results of the TLR7 branch-specific rate analyses (**Table 2**) suggest that the *Alouatta* clade is under significantly stronger purifying selection than the rest of the primates in the phylogenetic tree (*p* = 0.057). TLR7 in the Old World primate lineage evolves under significantly less stringent purifying selection (*p* = 2.0*e*^−05^) than the New World primates. TLR7 branch-specific rate analysis using only *A. palliata* also found that *Alouatta* has significantly lower *dN/dS* (*p* = 0.046). This indicates that the first TLR7 result is not due to using three *Alouatta* species and one species for each other genus. The TLR8 branch-specific rate analyses show no evidence that TLR8 *dN/dS* varies across the phylogenetic tree.

### 3.4 Branch site test and functional analysis

To identify specific codons subject to selection in the *Alouatta* lineage, we used a FEL test. Our null hypothesis was that the selection pressure for each codon is constant along the entire alignment. The alternative model allows the *dN/dS* rate to vary codon by codon along the alignment. For this test, significance is reached at 0.10 (Kosakovsky Pond and Frost, 2005). **Table 3** lists the codons evolving under a significantly different *dN/dS* rate in the *Alouatta* clade. FEL identified two sites under positive selection in TLR7 and six sites subject to positive selection in TLR8.

**Table 3:** Sites under positive selection in only in the howler monkey lineage identified by FEL analysis with functional implications.

| Gene | Codon Position | p-value | *Alouatta* Species | AA in *Alouatta* species* | AA other primates | Biochemical property difference (Y/N) | Functional Region |
| --- | --- | --- | --- | --- | --- | --- | --- |
| TLR7 | 538 | 0.048 | AGC | SER | PRO | Y | Binding |
| TLR7 | 721 | 0.097 | AGC | THR | ARG | Y | LRR |
| TLR8 | 800 | 0.042 | AGC, AC, AP | ALA | VAL | N | LRR |
| TLR8 | 248 | 0.055 | AGC, AC, AP | ILE | ILA *Siamiri*,<br>THR all others | Y | LRR |
| TLR8 | 284 | 0.063 | AGC, AC, AP | HIS | ASN | Y | LRR |
| TLR8 | 844 | 0.065 | AP | VAL | ALA | N | TIR |
|  |  |  |  |  | THR | Y |  |
| TLR8 | 211 | 0.071 | AGC, AC, AP | TYR | CYS/SER | Y | LRR |
| TLR8 | 744 | 0.097 | AGC, AC, AP | HIS | HIS *Callithrix*,<br>TYR/SER | Y | LRR |

Binding sites determined by (Wei et al., 2009). LRR regions determined by (Bell et al., 2003).
Abbreviations: AGC = *A. guariba clamitans*; AC = *A. caraya*; AP = *A. palliata*; AA = amino acid
\*As indicated in “*Alouatta* Species” column

Within TLR7, codons with lower *dN/dS* rates are due to a nonsynonymous variant in *A. guariba clamitans* (**Table 3**). The nonsynonymous change at TLR7 codon 538 is within the pathogen binding region and causes an amino acid change to serine (SER) only in *A. guariba clamitans*. Every other primate in the alignment has the amino acid proline (PRO) at TLR7 codon 538. Proline is non-polar while serine is polar. The second positively selected site is in the TLR7 LRR. At TLR7 codon 721 a nonsynonymous change produces the amino acid threonine (THR) in *A. guariba clamitans* while every other primate in the alignment has the amino acid arginine (ARG) at this site. Threonine and arginine are both polar, but threonine is neutral and arginine is positively charged. FEL analysis identified six variants under selection in functionally relevant regions of TLR8 (**Table 3**). Five of these variants are found in all three of the *Alouatta* species used in the study, and the last variant is unique to *A. palliata*. Five of the variants are within the LRR region of TLR8. The sixth variant is within the TIR domain.

## 4 Discussion

Previous studies evaluating evolutionary patterns of endosomal TLRs to identify species-specific differences have omitted the genus *Alouatta* in analyses (Quach et al., 2013; Wlasiuk and Nachman, 2010). Our analyses of TLR7 and TLR8 DNA sequences in primates revealed strong purifying selection on TLR7 in the *Alouatta* clade and a number of unique sites under positive selection in both genes. This study provides the first insight into genetic factors that may contribute to *Alouatta* susceptibility to YFV.

We first tested TLR7 and TLR8 sequences for disparity in branch-specific selection in *Alouatta* compared to other New World and Old World primates with variable susceptibility to YFV. While it is known that TLR7 and TLR8 are under strong purifying selection due to their essential role in virus recognition (Wlasiuk and Nachman, 2010; Areal et al., 2011; Rassa and Ross, 2003) our data showed that *Alouatta* TLR7 is subject to even stronger purifying selection than in other New World and Old World primate lineages. Since YFV originated in Africa and was introduced to the Americas by the slave trade recently, approximately 400 years ago (Bryant et al., 2007), *Alouatta* immune genes evolved under selection pressures from different ssRNA viruses. If the stringent purifying selection on TLR7 in the *Alouatta* lineage was necessary to maintain resistance to another virus, there would be little variation available to confer resistance to YFV, resulting in extreme susceptibility.

We also tested *Alouatta* TLR7 and TLR8 sequences for changes in codon-specific selection across the coding region. Despite strong purifying selection on TLR7, codon 528 in *A. guariba clamitans* is under positive selection. This site is in the pathogen binding region, meaning the amino acid directly interacts with ssRNA viruses, such as YFV, during infection (Wei et al., 2009). Amino acid differences within this region usually result from species-specific selection pressure (Quach et al., 2013; Wlasiuk and Nachman, 2010). The second positively-selected codon in TLR7 (codon 721) is due to an amino acid difference in *A. guariba clamitans* within the extra-compartmental LRR region. While the codon is not in the direct binding region, the LRR also plays a role in pathogen sensing and stabilization during the recognition process (Wei et al., 2009). Amino acid variations within TLR genes have been noted to change progression of a range of diseases in humans (Skevaki et al., 2015). Since both amino acid changes within TLR7 result in biochemical property changes at protein regions that interact with the pathogen, they are likely to affect binding or stabilization of the pathogen, potentially affecting YFV progression (Skevaki et al., 2015). Both amino acid differences in TLR7 are unique to *A. guariba clamitans*. The TLR7 phylogenetic tree reflects the FEL finding of unique non-synonymous changes in *A. guariba clamitans*, as the branch length is much longer than the other two *Alouatta* species due to a higher rate of substitution.

Future study of TLR7 in *A. guariba clamitans* individuals will be particularly important since the both positively selected variants identified in TLR7 are unique to the species. Additionally, the positively selected variants exist in spite of the resistance of TLR7 to substitutions in the *Alouatta* genus as a whole. *A. guariba clamitans* inhabits a region of Argentina with recent exposure to YFV due to deforestation (Di Bitetti et al., 1994; Holzmann et al., 2010). In 2007-2009, there was a YFV outbreak among the howler monkeys that resulted in mass deaths (Moreno et al., 2013; Holzmann et al., 2010; Bicca-marques and Freitas, 2010). Moving forward, it is imperative to collect samples from YFV affected *Alouatta* individuals to further explore whether the amino acid changes in *A. guariba clamitans* result in intragenus differences in YFV susceptibility previously unknown.

FEL identified six positively selected codons in TLR8. This is consistent with results from the branch-specific selection analysis, which found that TLR8 has been under less stringent purifying selection than TLR7 in *Alouatta* allowing more variation to arise in the gene. Five of the positively-selected codons in TLR8 are within the LRR region and four substitutions result in a biochemical change. Unlike the positively selected sites in TLR7, five positively selected sites in TLR8 are due to an amino acid difference in all three *Alouatta* species. The LRR region substitutions that cause a biochemical change could result in altered YFV sensing and immune system signaling in the *Alouatta* genus, affecting susceptibility. These five amino acid differences in TLR8 are found in all members of the *Alouatta* genus so it is likely that they arose earlier in *Alouatta* evolution before radiation and speciation across Central and South America. The sixth site, TLR7 codon 844, shows a nonsynonymous variant only in *A. palliata*. This site is within the TIR domain of the TLR. The TIR domain is the inter-compartmental region of a TLR and is responsible for initiating the signaling cascade to activate the innate and adaptive immune systems upon infection (Mikami et al., 2012). Species-specific differences in the TIR region are much rarer than in other coding regions (Areal et al., 2011; Mikami et al., 2012). Thus, the finding of a positively selected variant in the more highly conserved region of TLR8 warrants further investigation.

Typically, a complex combination of environmental, pathogen and host genetic factors play a role in determining susceptibility to infection. Our results reveal a number of genetic factors that could impact YFV susceptibility in *Alouatta*. It is essential to collect additional samples from *A. caraya* and *A. guariba clamitans* in YFV endemic ranges to clarify whether amino acid differences seen in this study are associated with YFV susceptibility. Moreover, further study may help identify genetic factors that influence susceptibility to YFV in humans and other primates.

## 5 Acknowledgements

We would like to thank Andy Burrell, Christina Bergey, and Todd Disotell for allowing us to use their *A. palliata* reference genome. We would like to thank the Brian Dalley at the University of Utah Huntsman Cancer Institute High-Throughput Genomics Center and Mike Powers and Derek Warner at the University of Utah Sequencing Core for assistance with sequencing protocols. We would like to acknowledge Marcela Uhart, Hebe Ferreyra, and the curators at the Museo Argentina de Ciencias Naturales Bernardino Rivadavia for their contributions. Funding for this project was provided by the Center for Global Change and Sustainability at the University of Utah and the Wildlife Conservation Society.

## Supplement

**Figure S1:**
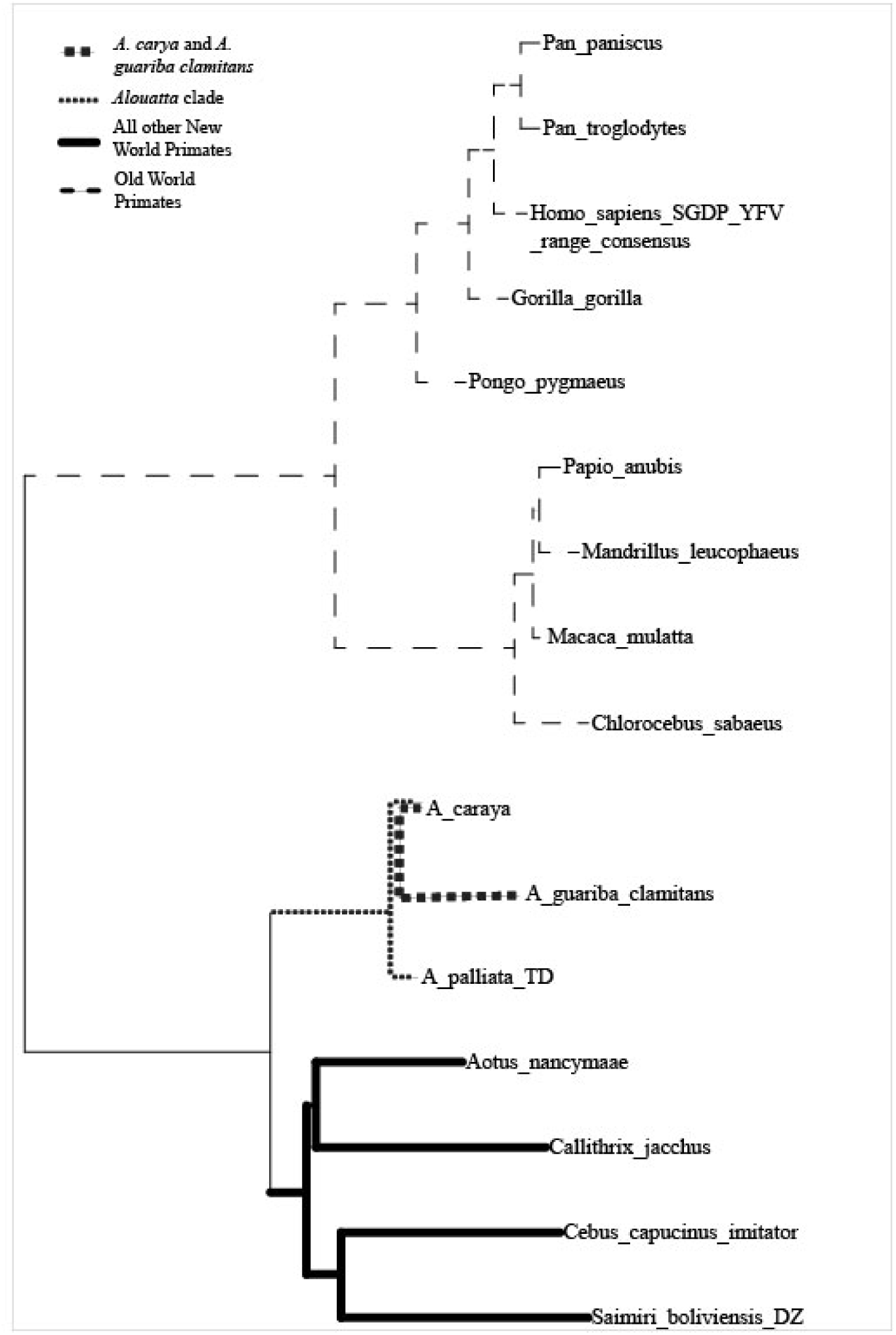
Illustration of branch bipartitions selected for SelectionLRT and FEL analyses.

**Figure S2:**
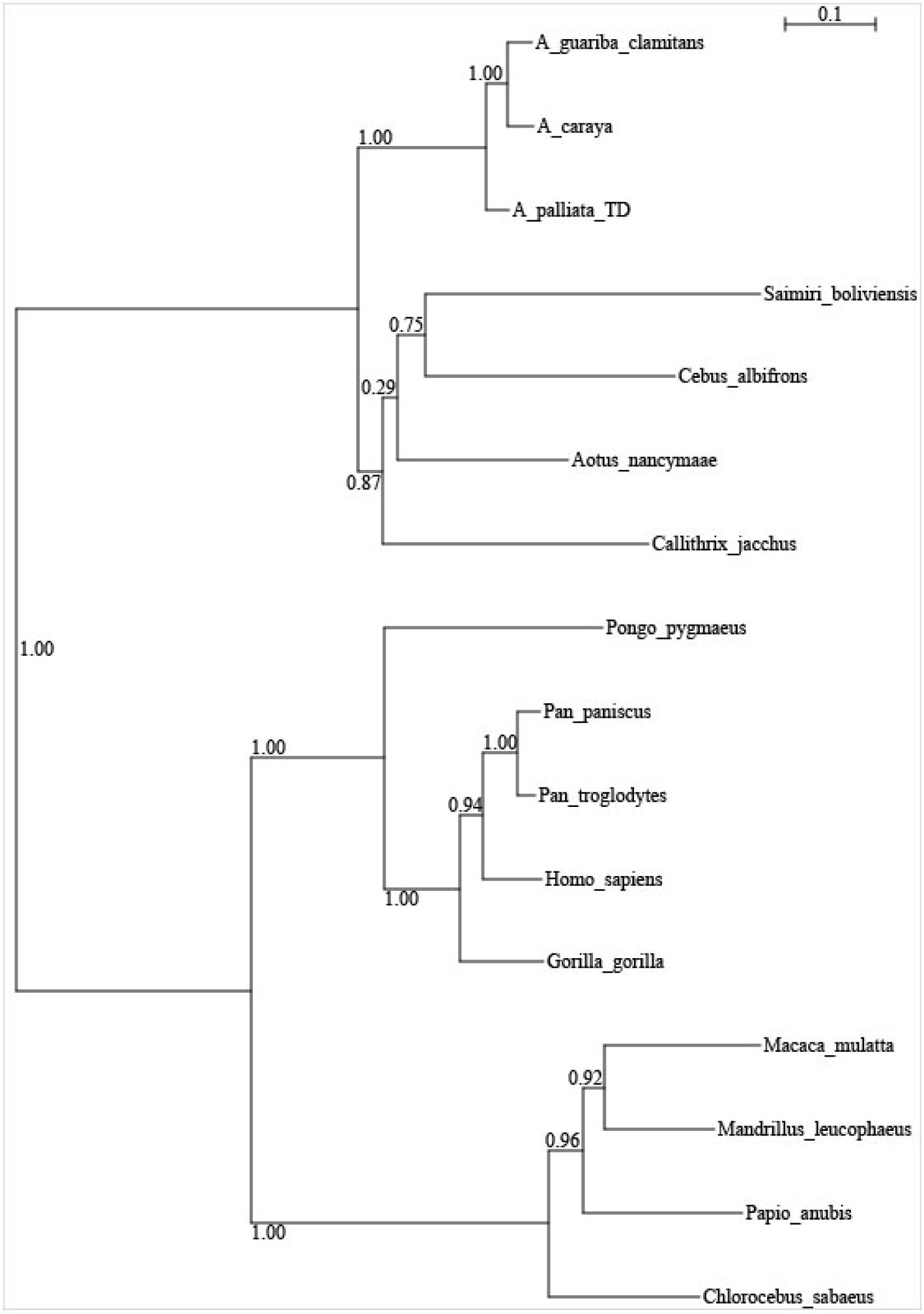
ND1-ND2-CO1 maximum likelihood phylogenetic tree generated from A. caraya and A. guariba clamitans TLR8 sequences (this study), New World and Old World primate mtDNA sequences **Table S3**. HKY model used, chosen by Decision Theoretic method. Each sequence totaled 3541 base pairs. Log likelihood of the tree = -24404.7 and aLRT support values on branches.

**Table S1:** Sample metadata

| Sample | Species | Collection location | Collection date | Sample type | Obtained from | Source ID | Sex |
| --- | --- | --- | --- | --- | --- | --- | --- |
| AC_tissue | *A. caraya* | Pindapoy Arroyo, San Jose | 2009 | liver tissue | Hernan Argibay | AC10 | M |
| AC_Museum | *A. caraya* | Corrientes, Argentina | 1967 | taxidermied skin | Museo Argentina de Ciencias<br>Naturales Bernardino Rivadavia | 13901 | F |
| AGC_Museum | *A. guariba clamitans* | Misiones, Argentina | 1952 | taxidermied skin | Museo Argentina de Ciencias<br>Naturales Bernardino Rivadavia | 52.41 | M |

**Table S2:**
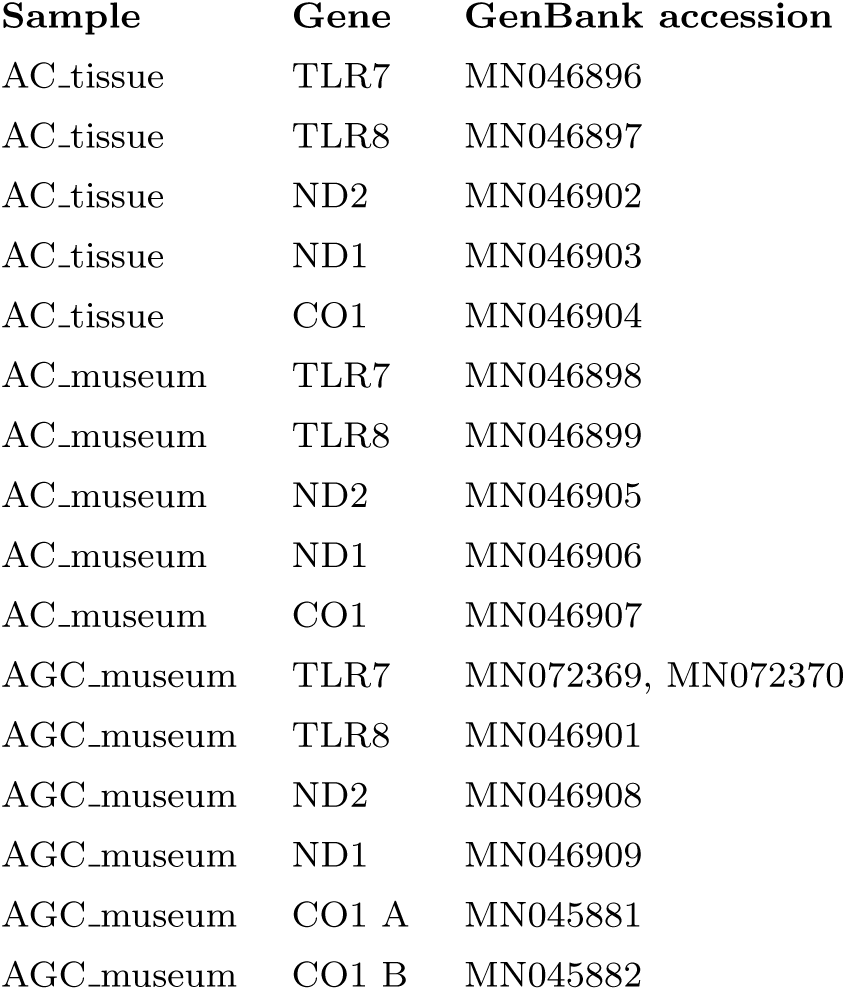
GenBank Accession Numbers

| Sample | Gene | GenBank accession |
| --- | --- | --- |
| AC_tissue | TLR7 | MN046896 |
| AC_tissue | TLR8 | MN046897 |
| AC_tissue | ND2 | MN046902 |
| AC_tissue | ND1 | MN046903 |
| AC_tissue | CO1 | MN046904 |
| AC_museum | TLR7 | MN046898 |
| AC_museum | TLR8 | MN046899 |
| AC_museum | ND2 | MN046905 |
| AC_museum | ND1 | MN046906 |
| AC_museum | CO1 | MN046907 |
| AGC_museum | TLR7 | MN072369, MN072370 |
| AGC_museum | TLR8 | MN046901 |
| AGC_museum | ND2 | MN046908 |
| AGC_museum | ND1 | MN046909 |
| AGC_museum | CO1 A | MN045881 |
| AGC_museum | CO1 B | MN045882 |

**Table S3:**
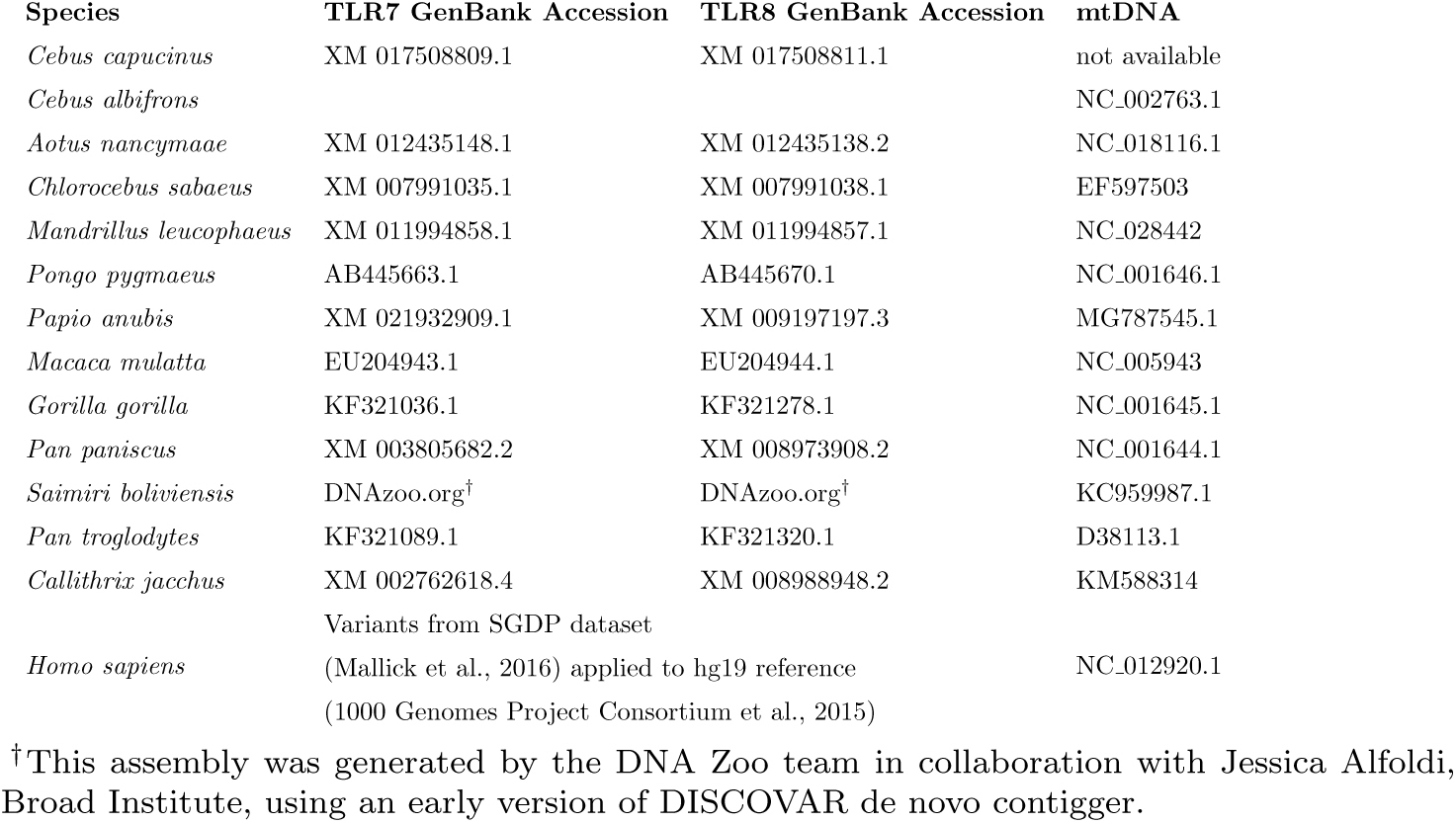
GenBank Accession numbers for TLR7 and TLR8 sequences used in phylogenetic analysis.

**Table S4:** Tree parameters: Phylogenetic tree parameters for each alignment as determined by Decision Theoretic method in PAUP (Swofford, 2003).

| Alignment | Model | Ts/Tv ratio | Invariable sites/<br>pinv | Across-site rate<br>variation/<br>Gamma rate |
| --- | --- | --- | --- | --- |
| TLR7 Coding Region | HKY | 4.54 | 0.747 | 0.171 |
| TLR8 Coding Region | TN | Optimized | 0.685 | 0.164 |
| mtDNA: CO1-ND1-ND2 | HKY | 27.33 | 0.391 | 0.32 |

**Table S5:**
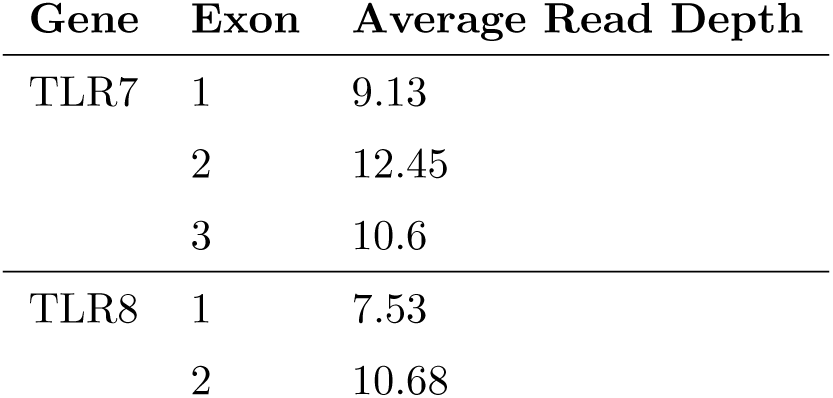
Read depth for at each gene and exon of *A. caraya* sample shotgun sequenced.

**Table S6:** Read depth for at each gene and exon of the *A. caraya* and *A. guariba clamitans* museum samples.

| Sample | Gene | Exon | Average Read Depth | Average Read Depth<br>after removing duplicates |
| --- | --- | --- | --- | --- |
| AC_Museum | TLR7 | 1 | 516.96 | 88.78 |
|  |  | 2 | 5050.54 | 289.12 |
|  |  | 3 | 1257.26 | 214.78 |
|  | TLR8 | 1 | 349.05 | 41.47 |
|  |  | 2 | 1037.97 | 182.96 |
| AGC_Museum | TLR7 | 1 | 21.94 | 6.42 |
|  |  | 2 | 80.83 | 9.07 |
|  |  | 3 | 43.98 | 8.33 |
|  | TLR8 | 1 | 10.51 | 3.94 |
|  |  | 2 | 60.63 | 9.74 |

**Table S7:** MapDamage2.0 results for museum samples.

| Sample | Output |
| --- | --- |
| AC_Museum | 0.00374 |
| AG_Museum | 0.00485 |

Values indicate the difference in C>T substitutions between the sample and the reference at the first 5' position. Values less than 0.01 indicate that level of DNA damage is low (Jónsson et al., 2013).

## Methods

We processed shotgun sequencing results and created a reference sequence using the following pipeline and commands:

1. Remove Illumina adapters from forward and reverse sequencing data (Bolger et al., 2014).

~~~
java −jar /bin/Trimmomatic 0.32/trimmomatic 0.32.jar
  PE threads 24 14457
 X1−sssss170823 D00294 0336 BCBLJ8ANXX 4 1.txt.gz 14457
 X1−170823 D00294 0336 BCBLJ8ANXX 4 2.txt.gz 14457
 X1−howlerX1 1P.fq.gz
 14457X1 howlerX1 1U.fq.gz 14457X1 howlerX1 2P.fq.gz 14457X1 howlerX1 2U.fq.gz
  ILLUMINACLIP:/usr/local/bwlab/bin/Trimmomatic
 −0.32/adapters/TruSeq3 −PE.fa:2:30:10:1:true
 LEADING:3 TRAILING:3 MAXINFO:50:0.2 MINLEN:50
~~~
2. Align forward and reverse howler monkey reads to a masked version of the human reference genome (hg19) (1000 Genomes Project Consortium et al., 2015) to reduce repetitive sequences (Li and Durbin, 2009). Output interlaced paired end read alignment.

~~~
 bwa mem −t 32 Homo sapien hg19/masked/hg19 masked
    14457X1 howlerX1 1P.fq.gz 14457X1 howlerX1 2P.fq.gz
~~~
3. Sort the aligned BAM files (Li, 2011).

~~~
 samtools view −bS hg19 masked 14457X1 howlerX1 PE.sam
   | samtools sort − hg19 masked 14457X1 howlerX1 PE.
   sorted.bam
~~~
4. BAM files filtered to target aligned reads to TLR7 and TLR8 and only include reads of quality MAPQ > 60 (Li, 2011).

~~~
 samtools view −q 60 −b {
   hg19 masked 14457X1 howlerX1 PE.sorted.bam}
   12885000−12941400 > hg19 TLR7−8 chrX
   :12885000−12941400 MAPQ60.fq
~~~
5. Create a consensus sequence of TLR7 and TLR8 using the fastq files (Bankevich et al., 2012).

~~~
 spades.py −−only−error−correction −t 24 −m 128
   −−12 hg19 TLR7−8 chrX:12885000 −12941400 MAPQ60.
   fq −o hg19 TLR7−8 chrX 12885000−12941400 MAPQ60
~~~

TLR7 and TLR8 are extremely conserved, especially within primates, therefore the howler monkey TLR7 and TLR8 genes were expected to align to the human reference genome. The same pipeline was used to align *Alouatta caraya* sequencing data to an *A. palliata* reference sequence provided by the A. Burrell at New York University. The final A. caraya TLR7 and TLR8 reference genomes created were identical.

We processed targeted sequencing results using the following pipeline:

1. Unalign BAM files for AC museum and AGC museum using GATK software (Van der Auwera et al., 2013).

~~~
java −jar /GATK Resource/picard−tools−1.134/picard.
  jar RevertSam I={input.bam} O={output.
   unaligned bam}
~~~
2. Convert unaligned BAM files to Fastq format (Quinlan, 2014).

~~~
bedtools bamtofastq −i {input.unaligned bam} −fq {
  output.fastq}
~~~
3. Assess fastq quality. Software available at http://www.bioinformatics.babraham.ac.uk/projects/fastqc/.

~~~
fastqc −o fastqc results sample.fastq
~~~
4. Trim fastq reads based on fastq quality results. Software available at https://sourceforge.net/projects/bbmap/.

~~~
bbduk.sh −Xmx1g in= {input.fq} out= {output.out fq} ftl
   =10 ftr=200 minlen=75 maq=20
~~~
5. Realign trimmed Fastq files for each sample to AC tissue reference (Li and Durbin, 2009).

~~~
bwa mem −R @RG*\\*tID:{id}*\\*tSM:{id}*\\*tLB:{id}*\\*tPU:{id}*\\*
   tPL: {IonPGM} {input.ref} {input.fq} | samtools
   fixmate −O bam − − | samtools sort −O bam −o {
   output.bams}
~~~
6. Index each BAM file (Li, 2011).

~~~
samtools index {input}
~~~
7. Mark duplicate reads using GATK software (Danecek et al., 2011).

~~~
 java −jar /GATK Resource/picard−tools−1.134/picard.
  jar MarkDuplicates I={input.bam} O={output.bam} M
  ={output.metrics}
~~~
8. Create a consensus VCF based on the BAM files (with duplicate reads marked) of each species (Garrison and Marth, 2012).

~~~
 freebayes −−fasta−reference AC tissue.fasta −−report
   −monomorphic −−genotype−qualities −−L {
   list of bam files} −v {output.vcf}
~~~
9. Remove indels from VCF file and filter by genotype quality (Danecek et al., 2011). Indels were removed since they appeared to be related to sequencing error caused by long strings of nucleotide Adenine.

~~~
 vcftools −−vcf {output.vcf} −−minGQ 30 −−remove−
   indels −−recode −out {output minGQ30.vcf}
~~~
10. Create a consensus sequence for the *A. caraya* and *A. guariba clamitans* taxidermied skin samples (Li et al., 2009).

~~~
 bcftools consensus −f ACref.fasta −s {sample} −H A −
  I output minGQ30.vcf.gz > sample consensus.fasta
~~~

